# Surface Plasmon Resonance Imaging of Excitable Cells

**DOI:** 10.1101/390948

**Authors:** Carmel L. Howe, Kevin F. Webb, Sidahmed A. Abayzeed, David J. Anderson, Chris Denning, Noah A. Russell

## Abstract

Surface plasmons are highly sensitive to refractive index variations adjacent to the surface. This sensitivity has been exploited successfully for chemical and biological assays. In these systems, a surface plasmon resonance (SPR)-based sensor detects temporal variations in the refractive index at a point. SPR has also been used in imaging systems where the spatial variations of refractive index in the sample provide the contrast mechanism. A high numerical aperture objective lens has been used to design SPR microscopy systems with the ability to image adherent live cells. Addressing research questions in cell physiology and pharmacology often requires the development of a multimodal microscope where complementary information can be obtained.

In this paper, we present the development of a multimodal microscope that combines surface plasmon resonance imaging with a number of additional imaging modalities including bright-field, epi-fluorescence, total internal reflection microscopy (TIRM) and SPR fluorescence microscopy. We used a high numerical aperture objective lens to achieve SPR and TIR microscopy with the ability to image adherent live cells non-invasively. The platform has been used to image live cell cultures demonstrating both fluorescent and label-free techniques. The SPR and TIR imaging systems feature a wide field of view (300 µm) that allows measurements from multiple cells while the resolution is sufficient to image fine cellular processes. The ability of the platform to perform label-free functional imaging of living cell was demonstrated by imaging the spatial variations in contraction of stem cell-derived cardiomyocytes. This technique has a promise for non-invasive imaging of the development of cultured cells over very long periods of time.

## 1. Introduction

Electrically excitable cells, such as cardiomyocytes and neurons, have been shown to produce fast optical signals that are the result of light scattering and birefringence changes associated with membrane depolarization [1–3]. In order to develop an understanding of how a population of cells are organised, exchange and process information a new sensing technology is required, one that has single cell resolution and can be used over a relatively large whole network. Surface Plasmon Resonance (SPR) sensors possess highly sensitive resonance conditions, which make them capable of detecting this membrane localized refractive index change with a high spatiotemporal resolution, label-free which will allow long-term recording.

SPR occurs when *p*-polarized light incident on a metal, at a specific angle of incidence, couples to the free electrons in the metal and all the incident energy is coupled into surfaces plasmons (SPs) [4]. At the angle of incidence where the plasmon coupling occurs, a drop in the intensity of the reflected light is observed and an evanescent wave is generated in both the metal and dielectric (Figure 1a). The generated evanescent field penetrates into the dielectric to a depth of about 150 nm, depending on the properties of the metal used [5,6]. SPR is very sensitive to perturbations of refractive index within this evanescent field; therefore, when there is a change in the refractive index of the dielectric medium, the characteristics of the light wave coupled to the surface plasmon changes and the resonance conditions are altered (Figure 1b). This allows SPR to be used for label-free imaging of features both structural (sensitive to local variations in density) [7–10], and functional (sensitive to movements of matter) [11–14].

**Figure 1.**
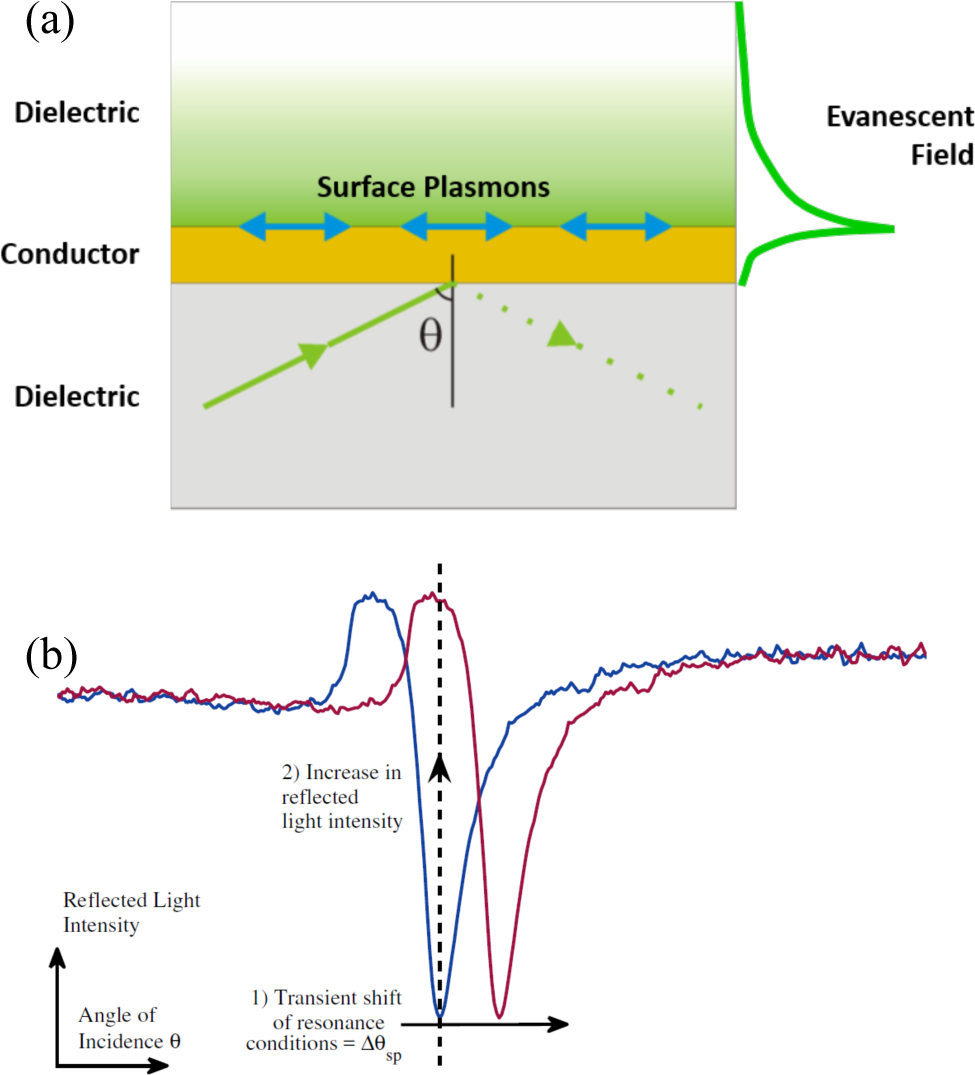
(a) Surface Plasmon Resonance on a conductive layer. When resonance occurs surface plasmons propagate along the interface between a conductor and dielectric. An evanescent field is generated that decays into both. (b) Diagram of angular and intensity modulation SPR sensor modalities. In an angular modulation SPR sensor the change in resonance conditions, for example, the refractive index of dielectric is detected by monitoring the change in the resonance angle, θsp. In an intensity modulation scheme, the angle of incidence is fixed and when θsp changes the detector sees an increase or decrease in light intensity.

Surface plasmons cannot be excited by the incident light directly. At a given photon energy (*h*ω) the wave vector of the incident *p*-polarized light must be increased so the photons can be coupled into plasmons [15]. Passing the illumination light through a high refractive index dielectric material, such as glass, can shift the wave-vector of the incident *p*-polarized light to excite surface plasmons. Excitation methods based on attenuated total reflection were demonstrated in the late sixties by Otto [16] and Kretschmann and Raether [17]. The Otto configuration uses a glass prism on top of a metal surface with a small air gap (~1 µm). However, the gap is difficult to control, so the Kretschmann-Raether configuration has a thin metal film deposited on top of a prism surface.

The Kretschmann-Raether configuration allows a large field of view (up to cm). However, this configuration is not suited to imaging because it requires a very shallow angle of incident light. The physical constraint of the prism limits the numerical aperture (NA) and magnification of the resulting imaging system and results in the configuration having a reduced spatial resolution [18]. Additionally, the field of view is angled relative to the objective resulting in an anisotropic distortion of the field of view. Using a high NA objective lens instead of a prism overcomes the issues of spatial distortion by keeping the object plane parallel to the image plane [18,19]. This method also allows higher magnification and, because of the high NA, the resolution of the imaging is much improved (diffraction limited: ~300 nm).

Despite these caveats, using SPR for functional imaging by detecting refractive index changes within the evanescent field is possible. To detect the change in refractive index of the dielectric medium interfaced to the metal surface, a number of detection schemes can be employed: (i) by tracking the minimum of the resonance angle (angular modulation) [20]; (ii) by monitoring the intensity at the highest gradient of the SPR curve (intensity modulation) [21,22] or using differential intensity detection approaches [23,24] or (iii) by detecting the phase of the reflected light [25]. The detection of small refractive index changes over a relatively large volume has been successful on some sensors based on an intensity modulation scheme down to a sensitivity (Δ*n_min_*) of 10^−6^ Refractive Index Units (RIUs) [21,22]. Better sensitivity levels have been achieved using other detection methods down to 10^−7^ RIUs using angular modulation [20]. Applying SPR imaging to research questions in cell physiology and pharmacology requires the development of a multi-modal system where complementary information can be obtained.

In this paper, we present a multimodal imaging platform that includes SPR microscopy using a high NA objective applied to live cell imaging with the following capabilities: (i) The SPR system features a wide field of view providing the ability to study ~40 cells simultaneously, with subcellular resolution. (ii) The SPR system is used to image neuronal cells while resolving axons and dendrites. We comment on the factors that affect the resolution of fine neuronal processes. (iii) We show that complementary information on the imaging resolution can be obtained from Total Internal Reflection Microscopy (TIRM) and the system, additionally, combines a number of microscopy systems that include bright-field, epi-fluorescence and SPR-excited fluorescence (SPRF) to be applied to cell physiology. (iv) The ability of the system to study spatiotemporal cellular functions is demonstrated by imaging localized contractions of stem cell-derived cardiomyocytes. (v) We describe a detailed design of this platform to enable ease of implementation, characterization for use by the cell physiology community.

## 2. Methods

### 2.1 Optical system design

The custom-built multimodal imaging system we have developed combines SPR imaging with a number of microscopy sub-systems that include bright-field, epi-fluorescence, total internal reflection microscopy (TIRM) and SPRF. Figure 2 shows a schematic of the optical system. The system is also capable of performing electrophysiological measurements simultaneously using microelectrode technology. The rig was mounted on a floating optical table (Thorlabs, Newton, NJ, USA) to minimize mechanical perturbations, to which such sensitive methods are highly susceptible.

**Figure 2.**
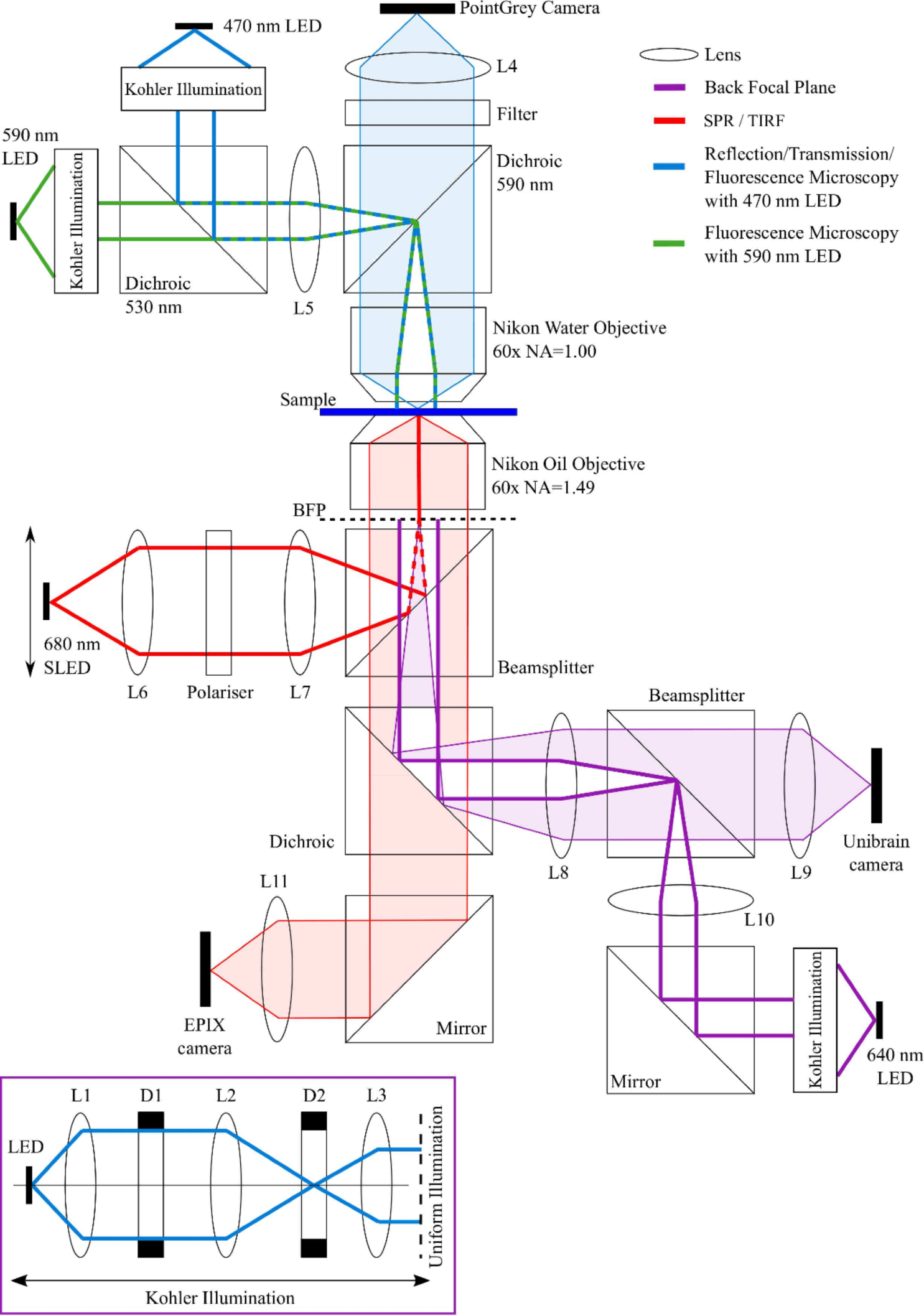
Schematic of the SPM that was developed. Several imagining modalities can be simultaneously exploited - including bright-field, epi-fluorescence, TIRFM, and SPR imaging Each optical pathway has been diagrammed within the figure. Imaging pathways are shaded, while the illumination pathways are shown with bold lines. Electrophysiology micro-manipulators (not shown) are also incorporated to provide for validation and calibration of the SPR signal with intracellular electrical activity.

A 640 nm light emitting diode (LED) was used in conjunction with Kohler illumination to illuminate the back focal plane (BFP) of the Nikon Oil Objective lens (CFI Apo TIRF 60×, NA = 1.49, oil-immersion lens, Nikon Inc, Tokyo, Japan) with a relatively uniform light intensity [26]. The BFP was monitored on the Unibrain camera (CMOS camera, 640×480 pixels, 5.6 µm pixel size, Fire-i^TM^, Unibrain Inc., San Ramon, CA, USA), shown in Figure 3a.

**Figure 3.**
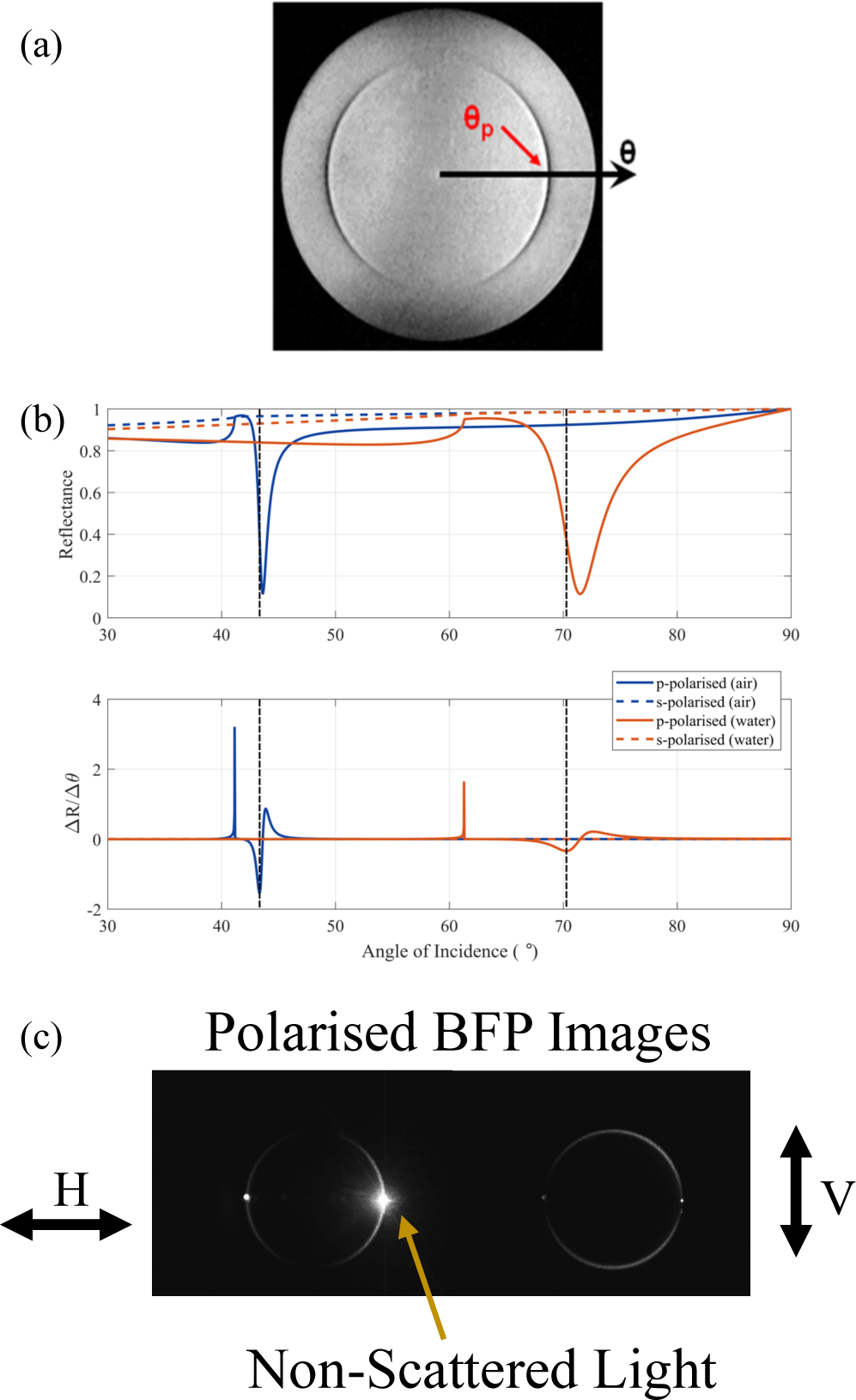
(a) Image of a uniformly illuminated back focal plane, with the gold sample in air. The dark arcs indicate the angles at which plasmon resonance occur. Note that only p-polarized light can excite plasmons, which is why arcs appear and not a ring. (b) Theoretical angular SPR response for p- and s-polarized light with air and water as the dielectric. S-polarized light is not capable of exciting surface plasmons. Using p-polarized light and increasing the refractive index of the dielectric increases the angle where resonance occurs. The angle where the gradient (ΔR/Δθ) is greatest is marked on each curve. (c) Images of the back focal plane imaged through a polariser with cells growing on a gold SPR surface. Note that light scattered off refractive index discontinuities in the sample returns from the same elevation angle but all azimuthal angles. This illustrates that momentum is conserved during plasmon scattering.

Reflection microscopy was performed with the 470 nm LED uniformly illuminating the BFP of a water-dipping objective (60×, NA = 1.00, Nikon Inc, Tokyo, Japan). The reflected light passed through the same objective lens, and was imaged on to Point Grey “Grasshopper3” monochrome camera (CCD camera, 1920×1440 pixels, 4.54 µm pixel size, S3-U3-28S5M-C Point Grey, Richmond, BC, Canada). Transmission microscopy was performed using the 470 nm LED, with the transmitted light imaged on the EPIX camera (CMOS camera, 640×480 pixels, 9.9 µm pixel size, SV643M, EPIX Inc., Buffalo Grove, IL, USA). The 470 nm and 590 nm LEDs could also be used for exciting fluorescence, in combination with appropriate emission filters.

Total internal reflection microscopy was achieved using the 640 nm LED, in conjunction with Köhler illumination to uniformly illuminate the BFP of the objective. The aperture diaphragm (D1) within the Köhler illumination was closed so that contrast was obtained in reflection by frustration of the critical angle (see supplementary information for information on the critical angle). The reflected light was imaged using the Unibrain camera.

When the objective pupil is filled with plane-polarized light from the 640 nm LED (Figure 3a), in the sector where the azimuthal angle is 0° the polarisation state is pure *p*-polarized light, while pure *s*-polarized light arrives at the sample where the azimuthal angle is ±90°. The dark arcs visible in Figure 3a show the angle of incidence where the light is coupled into surface plasmons and is therefore no longer being reflected. There is no dark band in the vertical direction (±90°) due to the light being pure *s*-polarized, and therefore incapable of exciting SPs.

SPR Fluorescence microscopy was performed by diffusional loading of Alexa Fluor®, 680-dextran (3kDa, Invitrogen™ D34681) into the cytoplasm of the cell of interest, using a micropipette attached to the cell soma. The fluorescent dye was then excited using the 680 nm superluminescent LED (SLED, 1 mW, Superlum Diodes Ltd, Cork, Ireland) tuned to excite plasmons beneath the cell of interest. The SPR-excited fluorescence was imaged on the and Point-grey Grasshopper3 camera. SPRF microscopy could be performed to confirm that the cells on the sample were in the evanescent field for SPR imaging.

The custom-built multi-modal SPR imaging system was developed around a high NA (NA = 1.49) objective lens and is able to exploit both angular and intensity modulation by either shifting the illumination angle (laterally translating the focal spot in the BFP) or holding this angle constant at the angle of maximum SPR gradient and monitoring the resulting intensity.

Angular modulation uses monochromatic light to excite the SPs, and the excitation of the SPs can be seen as a dip in the angular spectrum. The detector senses a shift in the angle where the reflected light intensity is at a minimum. Angular modulation was achieved using the 680 nm SLED, generating light that is focused into the objective BFP, resulting in an illuminating beam at the sample of an adjustable, narrow range of angles. A SLED was used over a laser or LED because they have a high spatial coherence but a low temporal coherence, allowing the output to be focused to a very tight spot [27,28], without suffering from the effects of laser speckling [29]. Additionally, SLEDs have extremely low noise [30], which is advantageous in the high-speed acquisition of rapid signals such as electrical activity in cells.

The output of the fiber-coupled 680 nm SLED was collimated using an achromatic lens and passed through a polarizer so that only *p*-polarized light was incident on the sample. The polarized SLED was focused to a diffraction-limited spot on the BFP of the objective lens with a second achromat, Figure 3c. The diffraction-limited spot, along with the polarizer, reduces the level of background caused by the *s*-polarized light that is unable to excite surface plasmons. The radial position of the illumination spot in the BFP dictates the angle of incidence at the sample. The light scattered off refractive index discontinuities in the sample when cells are cultured on the gold coverslips returns from the same elevation angle but all azimuthal angles. This illustrates that momentum is conserved during plasmon scattering. To change the angle of incidence at the sample the focused SLED beam was scanned across the BFP of the sample using a stepper motor traveling at 0.1 mm/sec. The reflected light was imaged on the EPIX camera at 24 fps, with an exposure time of 0.124 msec to provide reflection values as a function of the angle of incidence. Regions of interest were selected on the surfaces using Image-J and the *z*-axis profile was exported to MATLAB (Figure 3b).

Intensity modulation (IM) SPR sensing works by fixing the angle of incidence and wavelength and measuring the strength of the coupling between the light wave and the SP; detection is achieved by measuring the change in the intensity of the reflected light (Figure 1b). Therefore, for IM detection the angle of incidence (θ) was fixed at the angle with the steepest gradient, ΔR/Δθ (around 30% of the minimum intensity recorded at the trough of the SPR dip, marked on Figure 3b) and the variation of the reflected light intensity was monitored on the EPIX camera.

### 2.2 SPR Sensor Preparation

The glass coverslips (19 mm, Karl Hecht GmbH and Co KG, Sondheim, Germany) were treated with (3-mercaptopropyl) tri-methoxysilane (MPTS, Sigma Aldrich, 174617) to present thiol groups prior to thermal metal evaporation [31]. MPTS was used instead of the usual Chromium or Titanium adhesion layer to preserve the surface plasmon quality [32]. Silanisation was performed immediately after solvent and plasma cleaning. The MPTS deposition solution was prepared by mixing 1% v/v MPTS solution in anhydrous toluene. Upon removal from the plasma oven, the coverslips and rack were immersed in pure toluene for 1 minute. The rack and coverslips were transferred to the MPTS solution and left at room temperature for at least 30 minutes. Following immersion, the coverslips were rinsed in pure toluene, isopropyl alcohol, dried with a nitrogen line, then left to bake on a hotplate at 100°C for at least one hour - ideally overnight. The MPTS glass coverslips were coated with ~50 nm thick gold by thermal metal evaporation. Gold evaporation was carried out using a thermal evaporator (E306A, Edwards, Burgess Hill, UK). Gold wire was placed in a filament and the chamber was pumped down to 10^−1^ mbar with a rotary pump, then to 10^−7^ mbar with an oil diffusion pump, while being cooled with liquid nitrogen. A current of ~30 A was passed through the filament, so the metal melted and evaporated onto the glass coverslips held above the source with rare earth magnets. An Intellemetrics IL100 quartz crystal microbalance was used to estimate the evaporation rate and film thickness. The thickness of the gold coverslips was measured with a spectroscopic ellipsometer (Alpha-SE, J.A. Woollam, Lincoln, NE, USA) at a fixed incident angle (70°) and with a wavelength range between 380 - 900 nm.

### 2.3 Sensitivity Experiments

Sensitivity experiments were performed on the planar gold surfaces to determine the minimum refractive index change (sensitivity) that could be detected. Different concentrations of Sodium Chloride (NaCl) salt solutions in purified water were added to a chamber on top of a gold-coated coverslip whilst the reflected SPR light intensity response was measured. Deionized water was added as the first sample to find the reflection gradient maximum and the SLED was positioned here for the entirety of the experiment. While keeping all other parameters fixed, the refractive index of the media was adjusted using different concentrations of salt solution. The change in refractive index was calibrated with an Abbe Refractometer (Abbemat 200, Anton Paar, Graz, Austria). The reflected light from the sample plane was recorded using the EPIX camera at ~286 Hz. To allow the solution to stabilize, 30 – 60 seconds was left between each step. The resulting intensities were exported using Image-J and further analysis was performed in MATLAB.

### 2.4 Substrate Preparation

For cell-based experiments on glass, the coverslips were treated with poly-L-lysine (PLL, Sigma Aldrich, P4707). The PLL coated glass coverslips were prepared by immersion in PLL solution for 5 min followed by thorough rinsing with deionized water.

For cell-based experiments on planar gold, the coverslips were treated with 11-Amino-1-undecanethiol (AUT, Sigma Aldrich, 674397), to generate a self-assembled monolayer [33]. First, the gold coverslips were solvent cleaned, followed by an oxygen plasma treatment.

The AUT was dissolved in ACS grade Ethanol at 1 mM, which was then poured into an immersion cylinder to completely cover the gold. Immersion time was not less than 18 hours. Following immersion, the solution was drained from the cylinder and the gold was thoroughly rinsed, first with ethanol to remove the bulk AUT solution, and then with distilled water to remove the cytotoxic ethanol. The coverslips were dried thoroughly with N_2_ before cell plating.

### 2.5 Cell Culture

The 3T3 fibroblast cell line was maintained in Dulbecco’s Modified Eagle Medium (DMEM, Gibco, 11960) with 10% Fetal Bovine Serum (FBS, Gibco, 10270). Cells were subcultured and seeded on to AUT treated gold coverslips at a density of 0.3×10^6^ cells.

The stem cell-derived cardiomyocytes were differentiated on PLL coated glass coverslips from human embryonic stem cells (hESC-CM) following the protocol in [34].

Cultures of primary hippocampal neurons were dissected from E18 Wistar rat embryos following a standard protocol [35]. The primary rat hippocampal neurons were plated on the functionalized coverslips with ~150,000 dissociated cells in 500 µl media.

## 3. Structural Imaging

### 3.1 Optical System Characterisation

#### 3.1.1 Field of View

Functional imaging of excitable cells using SPR microscopy requires performing measurements from a number of single cells to inspect the cell-cell interactions and replicate single cell measurements. Therefore, the system has been designed with a wide-field of view to allow for imaging of multiple single cells with submicrosocpic resolution. This has been achieved by deliberately reducing the magnification of the system.

The field of view was increased by mismatching the objective and tube lens to reduce the effective magnification of the 60× oil-objective. The Nikon objective lens is designed for a tube lens with a focal length of 200 nm, however, we used a 60 mm tube lens. This reduces the effective magnification to 18× and enlarges the field of view to up to 500um, depending on the size of the camera sensor.

The number of pixels in the EPIX camera sensor is 640×480, at 9.9×9.9 µm each, so the field of view (FOV) has been increased to 350×250 µm from 105×80 µm. Figure 4 shows SPR images of mouse fibroblast cells (3T3) seeded on gold coverslips, showing the wide field of view and the potential to image relatively large populations of cells, for example, there are ~40 cells within Figure 4.

**Figure 4.**
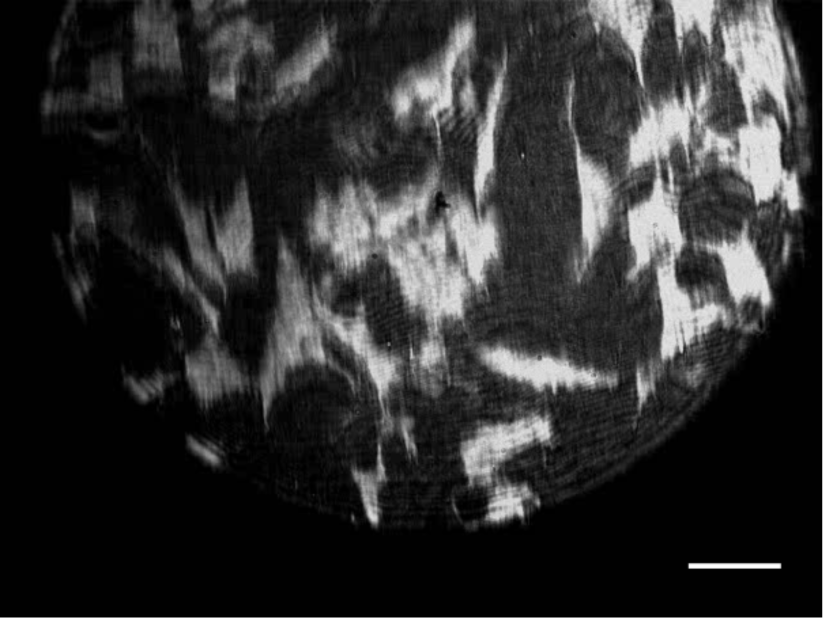
SPR image of cultured fibroblasts indicating high contrast over a wide field of view (350×250 µm). The scale bar is 40 µm.

#### 3.1.2 Contrast and Spatial Resolution

The contrast is limited only by the spatial coherence of the SLED. The resolution of a conventional light microscope is limited by the diffraction limit. However, in SPR sensors, the plasmons propagate along the metal/dielectric interface for a distance determined by the propagation length before they decay back into photons. This limits the resolution of the system in the direction parallel to the propagation of SP waves. Figure 5a shows a primary, hippocampal neuron cultured on a gold sensor imaged at 5 days *in vitro* (DIV). The fine, elaborating features visible in the Figure demonstrate that axons and dendrites can be clearly resolved using SPR imaging in this regime. However, cell processes that are perpendicular to the propagation of the SPR are hardly resolved, due to the resolution limit of the SPR system, while they are clearly visible when they are parallel to the propagation of the SP wave [36].

**Figure 5.**
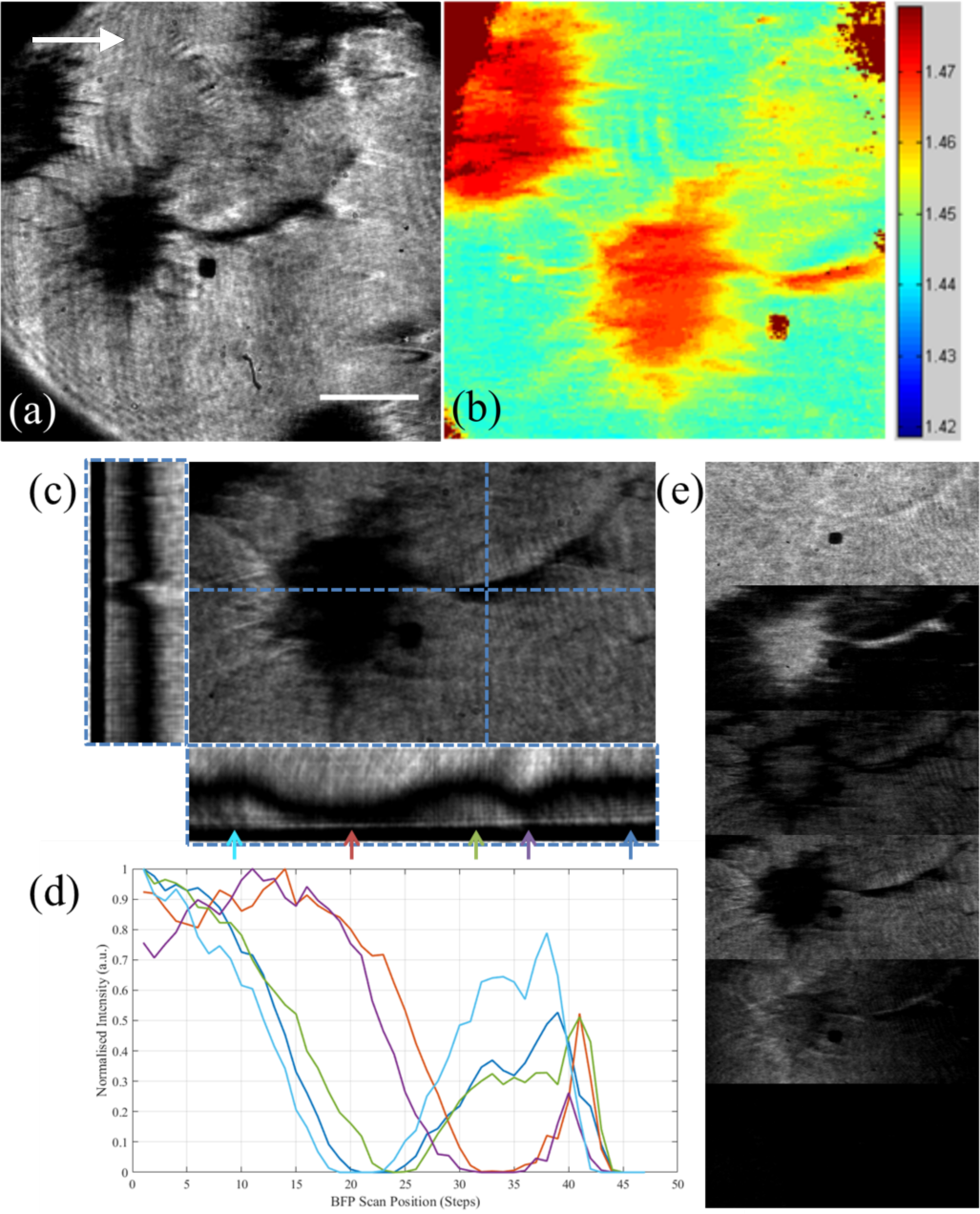
(a) SPR image of a neuron cultured on the SPR sensor. Arrow indicates the direction of plasmon propagation. (b) Pseudo-color image showing the effective refractive index (in RIU) at each pixel in the field of view. This was created by scanning the illumination angle and finding the angle of minimum reflection for each pixel. The effective refractive index is reasonably homogeneous underneath the soma indicating the cell has adhered to the surface uniformly. The scale bar in (a) is 10 µm long and consistent across both images. (c) BFP angle-scan, with vertical and horizontal projections of the stack – dark line represents the SPR dip, with the “Z” axis encoding the BFP angle. (d) Angle scan at various points across the cell and coverslip, extracted from the projection in (c), indicated by arrows. (e) Successive adjustments of SPR excitation – top to bottom: below SPR angle, SPR on a coverslip, middle of SPR dip on cell, SPR under cell, edge of BFP, outside BFP (dark).

In biological samples, spatial differences in the lipids and proteins present result in a variation of the refractive index. These differences provide the contrast in an SPR image. When the propagating plasmon reaches a refractive index boundary it is no longer supported and is re-radiated as photons. This means that, even though in theory the plasmon propogation length is typically 20 µm, fine details, such as neuronal axons and dendrites can be resolved. Note that the contrast is also dependent upon the distance betweeen the cell and substrate [8,10].

To assess the flatness of cell adhesion on our gold surfaces, the refractive index of each pixel in the field of view of Figure 5a was determined by scanning the illumination angle and finding the angle of the minimum reflection for each pixel, Figure 5b and Figure 5d. Then solving the Fresnel equations for reflection and transmission in a multi-layer device the refractive index was calculated, Figure 5b (see supplementary information). The effective refractive index is reasonably homogeneous underneath the soma indicating the cell has adhered to the surface uniformly. Reflection as a function of scan angle at one *x*-position and one *y*-position are shown in Figure 5c. The dark line in the projections represents the SPR dip and corresponds to the angle of incidence at the gold layer. The complete angle scan is shown at various points on the sample in Figure 5d, while Figure 5e shows the full image at various angles during the scan. Note that at different angles of incidence plasmons are excited sucessively either under the cells or on cell-free background areas.

The spatial resolution defines the smallest distance between two Airy disks that can be resolved [37]. The theoretical diffraction-limited resolution with the original tube lens for a 60×, 1.49 NA microscope objective for 680 nm incident light given by the Abbe limit is 0.28 µm. However, at a magnification of 18× the minimum resolvable feature, which is magnified onto a 2×2 region of the sensor in accordance with the Nyquist criteria, is 1.1×1.1 µm. The average measured size of a typical cultured mammalian neuronal soma is ~16 µm (data not shown) [38] and a typical cardiomyocyte cell is about 100 µm long and 10 – 25 µm in diameter [39]. This resolution is, therefore, more than adequate for imaging both cardiomyocytes and networks of neurons including many of the dendrite and axonal processes which are typically 1 – 2 µm in size [40].

To demonstrate the resolution limitations of the SPR microscopy, we explored an alternative evanescent wave microscopy technique that is based on total internal reflection (Figure 6). Primary hippocampal neurons were cultured on glass coverslips treated with PLL. The glass surface was illuminated with a uniform disc of light from the 640 nm LED source objective BFP. This disc of light was stopped down using the aperture diaphragm (D1) to just above the total internal reflection critical angle to gain contrast.

**Figure 6.**
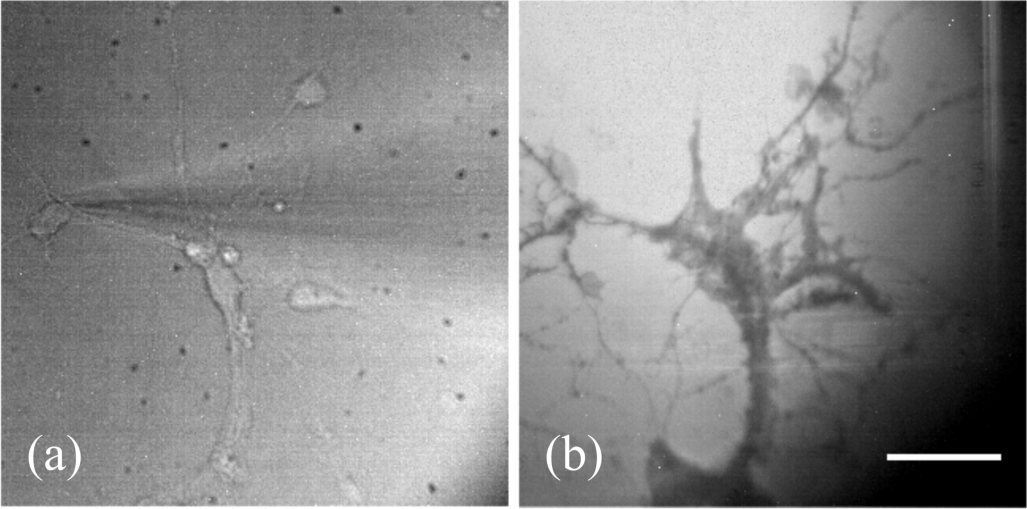
Transmission (a) and TIRM (b) images of primary hippocampal neurons cultured on glass surfaces. (a) The glass surface was illuminated with the 470 nm LED and imaged on the EPIX camera. The microelectrode is visible in the image. (b) The 640 nm LED uniformly illuminated the back focal plane of the objective. The light was stopped down to close to the TIR critical angle to gain TIRM contrast. The scale bar is 40 µm long and consistent across both images.

These results demonstrate that TIRM provides better resolution compared to SPR and simple transmission microscopy (Figure 6a). The results obtained were performed using a glass substrate, however, these can be achieved using gold-coated sensors to allow one to obtain complementary TIRM images with the same substrate that is used for SPR, by only changing the angle of incidence to 61.3° for an air/water interface. Although TIRM provides resolution and contrast based on the frustrated total internal reflection, it has a limited ability to resolve functional time-resolved information compared to SPR.

### 3.2 SPR-excited Fluorescence Imaging

A microelectrode filled with dye was patched into the cell to demonstrate that the neurons were close enough to the gold surface for their membranes to lie within the evanescent field. The dye within the cell was excited using both epi- and SPR-excited fluorescence, Figure 7a, and 7b, respectively. Figure 7b confirms that the 680 nm SLED SPR system can excite the dye-labeled cell and therefore, the cell is within the evanescent field. This is a necessary prerequisite to allow functional imaging using SPR. 100% of cells tested (N=12) were found to lie within the evanescent field. This shows that SPR could be used to excite calcium or voltage dyes in studies of neuronal (or other) cultures.

**Figure 7.**
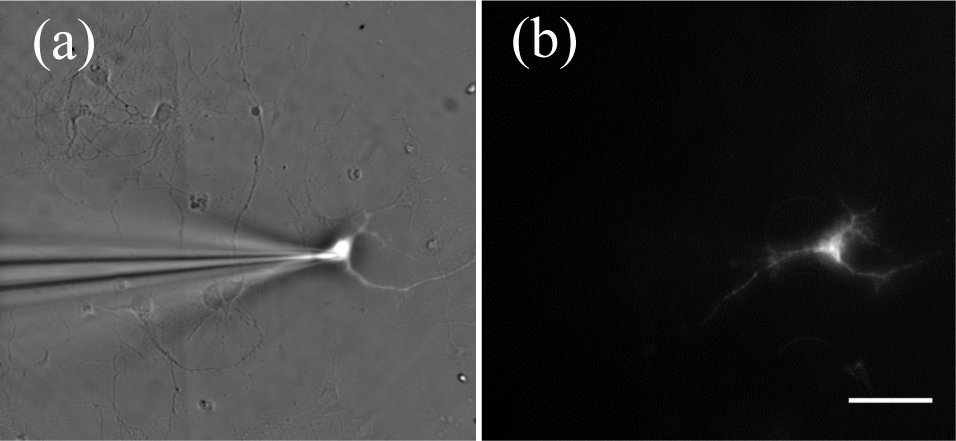
Images of a cultured neuron on an SPR sensor. (a) A microelectrode filled with a fluorescent dye is attached to a neuron imaged using reflection microscopy which, can be seen under epi-fluorescent illumination. (b) Neurons filled with fluorescent dye illuminated with the SLED with SPR. This confirms the neurons are closely adhered to the gold SPR surface and within the evanescent field.

## 4. Functional Imaging

For label-free functional imaging of dynamic cellular functions, from cardiomyocytes for example, a low noise, high sensitvity system is required. We characterized this sensitivity by changing the bulk refractive index of the dielectric. This characterisation is followed by results that demonstrate the ability of the system to image localized dynamic processes. As described later in section, the system has been used to image spatial variations in the contraction of stem cell-derived cardiomyocytes.

### 4.1 Sensitivity to refractive index changes

To characterize the sensitivity of the planar gold surfaces on the SPM, the minimum detectable change in refractive index was experimentally determined by changing the bulk refractive index of the dielectric on top of a 50 nm planar gold sample.

The concentration of the medium on the surface on the substrate was changed to increase or decrease the refractive index using NaCl solutions. The concentration values were chosen so that the change in refractive index in step sizes of Δ*n* = ~0.0001. To find the minimum detectable change (the sensitivity), the angle of incidence must be chosen such that the reflection gradient (ΔR) is at its maximum. Practically, it is difficult to set this parameter, so the dynamic range was first established by angle-scanning across the SPR dip, and then setting SPR illumination to the location of the greatest gradient (Figure 3b). There is a trade-off between greater sensitivity and SNR, as at the angle of the maximum gradient the light intensity is much lower, reducing the Poisson-limited SNR. Considering this, the angle of incidence was fixed at around 30% of the dynamic range. A 1 mW 680 nm SLED light source was used with the 12-bit CMOS camera and a 3.5 msec exposure time.

The results from all the experiments are summarized in Figure 8. Figure 8 shows that the normalized light intensity (Δ*R*) increases linearly with increasing refractive index. The sensitivity of the sensor was characterized by the refractive index unit (RIU). The sensitivity in RIU was calculated as 2×10^−5^.

**Figure 8.**
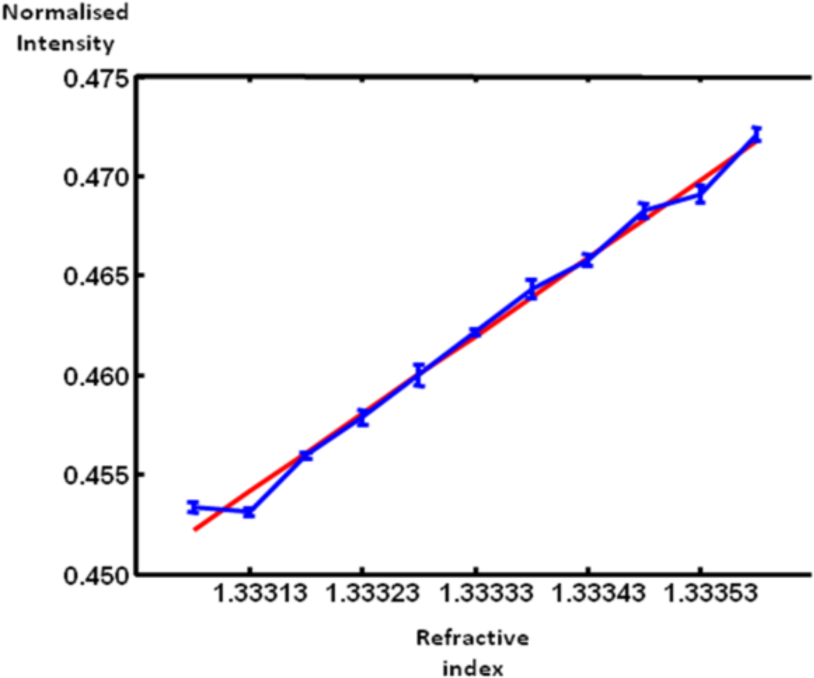
The sensitivity of the SPR imaging system. Refractive index was adjusted using a series of different concentration NaCl solutions. A 1 mW 680 nm SLED light source was used with a 12-bit CMOS camera and a 3.5 msec exposure time. The sensitivity in RIU was calculated as 2×10^−5^.

The experimentally measured Δ*n*_*min*_ here is less than is the values of ~10^−6^ RIU reported from intensity modulating SPR techniques in the literature [21,22]. However, the level of detection in these sensitivity experiments is limited by the exposure time because the power of the noise is proportional to the bandwidth. So for the small bandwidth used for these measurements (286 Hz), the total noise power may be reduced if a lower sample frequency is used and the SNR increased.

### 4.2 Detection of cardiomyocyte contraction

Stem cell-derived cardiomyocytes were cultured on gold surfaces. The periodic contractions of a living, beating cell could be visualized by monitoring the change in SPR intensity on the EPIX camera. By taking the difference of each subsequent frame with the first frame a stack was produced that shows the area where there is the greatest variation in light intensity (‘‘hot-spot”), Figure 9c and 9d. The background was corrected for by taking the difference of the ‘‘hot-spot” with a sample of the rest of the cell which was not moving.

**Figure 9.**
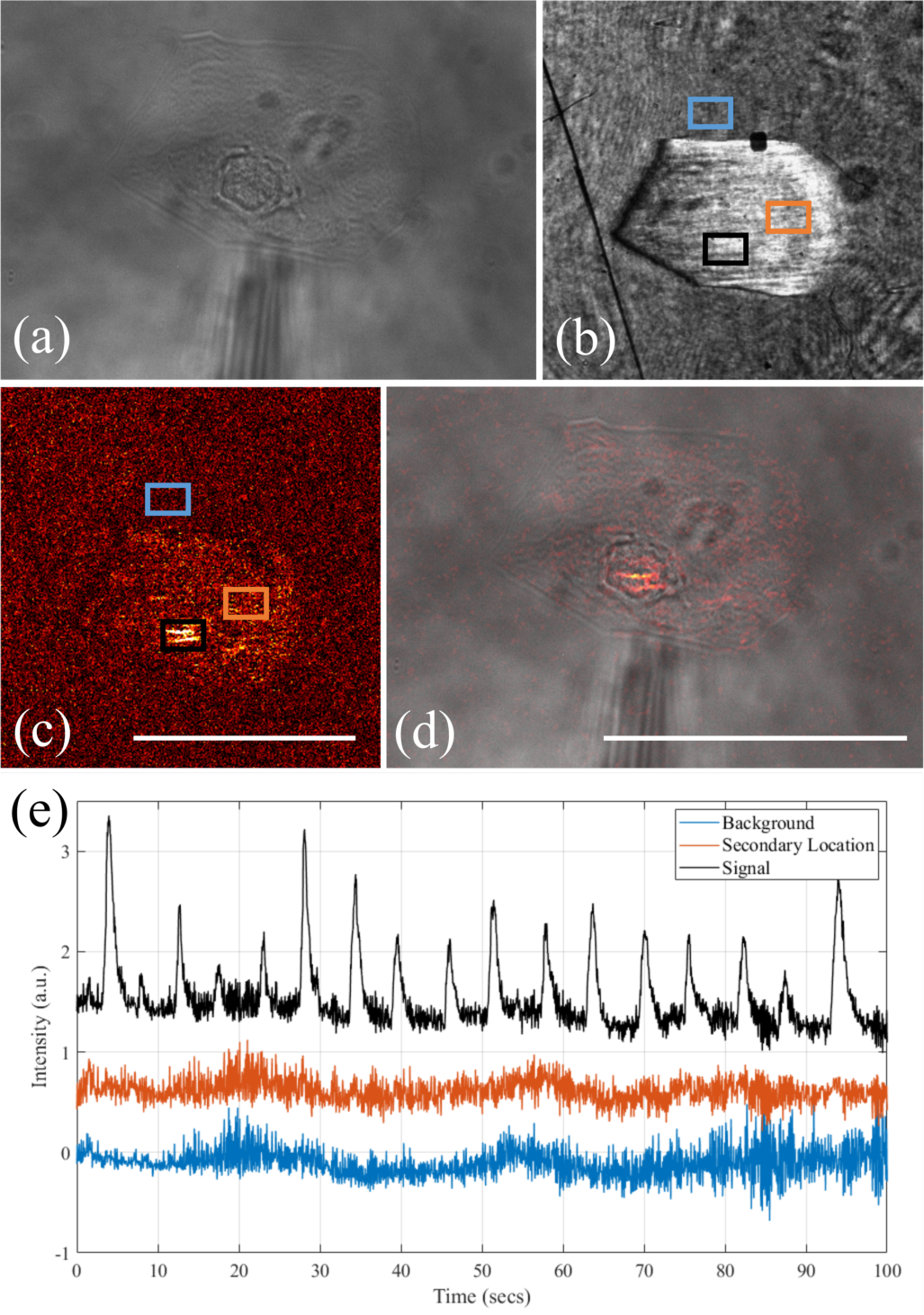
(a) Bright-field image of a stem cell-derived cardiomyocyte taken using reflection microscopy. (b) SPR image of the cell from (a). (c) There are two stripes of localized light intensity changes, which can be seen when subtracting the first frame of the SPR response from the remaining image stack. The scale bar is 10 µm long and consistent across (b) and (c). (d) Overlay of the (c) on (a). The area of the greatest light intensity change appears to be around the nucleus of the cell. The scale bar is 10 µm long and consistent across (a) and (d). (e) Relative change in the intensity of the SPR signal from a beating stem cell-derived cardiomyocyte, a control area directly next to the cell of interest and a second location within the cell. The ROIs are highlighted in (b) and (c). Each trace was offset for clarity.

Figure 9e shows that the SPR intensity changes in time with the contractions of the stem cell-derived cardiomyocyte (‘‘signal’’) with a high signal-to-noise ratio. Two control ROIs are plotted. One on the background and a secondary location within the cell. The resulting SPR responses shows that only a couple of areas are moving within the cell, with the rest being uniform. The area where there is the greatest change in light intensity appears to be around the nucleus of the cell. It is possible that the source of the signal is from the stress fibres or contractile apparatus within the cardiomyocyte pressing on the nucleus causing it to move. The control ROI on the ‘Background’ was taken directly next to the beating cardiomyocyte, which shows no change in relative SPR intensity. This demonstrates that the change in SPR intensity is localized to the cell of interest and therefore, most likely due to refractive index changes or localised movement within the cell and not from movement causing mechanical waves throughout the volume. A video of the time series is given in the supplementary information.

## Conclusions

In this paper, we have presented a multi-modal platform for cell physiology combining SPR imaging with a number of ancillary microscopy systems. We have shown that the system is capable of both structural and functional imaging of cultured cells. Using the system for structural imaging a number of modalities can be exploited including reflection/transmission microscopy, TIR micrscopy, epi-fluorescence and SPR imaging to obtain complementary information. A wide field-of-view has been demonstrated with a suitable spatial resolution for imaging cardiomyocytes and resolving individual axons and dendrites in cultured primary neurons label-free.

The ability of the system to study spatiotemporal cellular functions was demonstrated by imaging localised contractions of stem cell-derived cardiomyocytes. To the best of the author’s knowledge, this is the first demonstration of functional imaging of the refractive index changes with SPR in single cardiomyocytes. Using SPR could allow the localised contractions of cardiomyocytes to be imaged in real-time and drugs to be tested *in vitro*. In future work, a small network of cultured neurons will be grown on the surface of a sensor and any small changes to this light that occur during an action potential will be monitored.

## Acknowledgments

The authors would like to acknowledge financial support from The Engineering and Physical Sciences Research Council (EP/F005512/1 and EP/H022112/1).) and a Royal Academy of Engineering/EPSRC Post-doctoral Fellowship to K.F.W. (EP/G058121/1). Authors would also like to thank BHF (SP/15/9/31605, RG/15/6/31436, PG/14/59/31000, RG/14/1/30588, RM/13/30157, P47352/CRM); BIRAX (04BX14CDLG); MRC (MR/M017354/1); NC3Rs (CRACK-IT:35911-259146, NC/K000225/1). C.L.H would like to acknowledge financial support from The Whitworth Society.

